# Developing a versatile technology for agroinfiltration in multiple plants

**DOI:** 10.1101/2025.05.25.656011

**Authors:** Tiange Wang, Jieyu Ge, Chengyi Qu, Hongyu Chen, Jie Li, Shun Wang, Xueying Guan, Tao Chen, Zhanfeng Si, Keming Hu, Xinhua Ding, Li Liu, Jian Zhang, Alexander S. Mishin, Ilia V. Yampolsky, Hao Du

## Abstract

*Agrobacterium*-mediated transient transformation methods are widely used in plant science and molecular pharming. However, plant immune response and post-transcriptional gene silencing (PTGS) can compromise gene expression, limiting the adaptability of these methods across plant species. The lack of a robust reporter system further hinders advancements in agroinfiltration technology. To address these challenges, we developed a fungal bioluminescence pathway (FBP)-based reporter system that utilizes caffeic acid to generate self-sustained luminescence. By incorporating NahG and P19, our FBP assay enhances transient transformation efficiency in over 20 plant families, including major crops, vegetables, ornamental plants, and trees. Furthermore, we demonstrate its utility for assessing protein localization, interactions, and transcriptional regulation in multiple plant species, thereby broadening our understanding of gene function throughout the plant kingdom.

## Introduction

Transient expression in plants is a cutting-edge approach that enables rapid, high-throughput evaluation of gene function, thereby accelerating functional genomic studies. In particular, agroinfiltrated *N.*LJ*benthamiana* can quickly yield large amounts of foreign proteins, making it an appealing system for molecular pharming (Beritza *et al*., 2024; Lomonossoff and D’Aoust, 2016). Compared with mammalian cell culture or microbial fermentation, plant-based transient expression provides unique advantages such as lower production costs, reduced risk of human pathogen contamination, scalability, and the ability to carry out complex post-translational modifications such as disulfide bridges and glycosylation (Murad *et al*., 2020; Tschofen *et al*., 2016). Despite these benefits, agroinfiltration-based transient expression remains challenging for many plant species, including crops, ornamental plants, and perennial trees. Although methods like biolistic bombardment and agrobacteria infiltration have been optimized for transient transformation in species such as *Arabidopsis*, *Solanum tuberosum*, and *Capsicum annuum* have been documented (Rosas-Díaz *et al*., 2017b; Ueki *et al*., 2009; Zhang *et al*., 2020b), their wider adoption is hindered by specialized equipment requirements, the expense of gold microparticles, and, notably, low transformation efficiency.

Plant resistance to *Agrobacterium* is closely linked to the activation of plant immune responses upon detection of infiltrated *Agrobacterium*, thereby inhibiting the transformation efficiency. Specific pathogen-associated molecular patterns (PAMP) from *Agrobacterium* can trigger a signaling cascade that leads to PAMP-triggered immunity (PTI) (Zipfel *et al*., 2006). However, this approach may not be universally applicable due to the mutant background of the plants. Previous studies have explored different methods to enhance transient transformation, such as utilizing transgenic plants expressing bacterial type-III effector AvrPto to inhibit immune-related kinases (Tsuda *et al*., 2012). In the context of the plant-*Agrobacterium* pathosystem, salicylic acid (SA) plays a key role in orchestrating defense reactions. Studies have shown that *Arabidopsis thaliana* and *N. bethamiana* plants with defective SA accumulation are more susceptible to infection (Anand *et al*., 2008; Yuan *et al*., 2007). Thereby, another promising approach involves the expression of the NahG, encoding bacterial salicylate hydroxylase, in *Arabidopsis*. This strategy has been found to significantly enhance the efficiency of *Agrobacterium*-mediated transformation in rosette leaves (Rosas-Díaz *et al*., 2017c), facilitating the routine use of this technique in agroinfiltration assays. However, challenges remain in extending the effectiveness of this method to a wider range of plant species.

Foreign transcripts in plants often become unstable due to PTGS, a process that typically begins three days after agroinfiltration. PTGS is initiated by the presence of low levels of antisense RNA, which can be produced by random transfer DNA insertion and/or RNA-dependent RNA polymerase. This leads to the formation of double-stranded RNA, which is then processed by Dicer into small interfering RNAs (siRNAs) that target and degrade homologous mRNAs (Gopalakrishnan and Wolff, 2009). One approach to counteracting PTGS is the removal of Dicer-like proteins (DCLs) DCL2 and DCL4, in the *dcl2dcl4* double mutant of *N. benthamiana*, which has been shown to increase transient expression of recombinant proteins (Matsuo and Matsumura, 2017). In the ongoing battle between hosts and pathogens, viruses have developed viral suppressors of RNA silencing (VSRs) as a means of evading host defense mechanisms (Wu *et al*., 2010). Out of all the VSRs, the plant tombusviral P19 protein is notably one of the most thoroughly researched at the biochemical level. It effectively suppresses PTGS by binding to and sequestering siRNAs (Lombardi *et al*., 2009). Despite its well-documented effects, the efficacy of P19 in agroinfiltration experiments across different plant species remains to be fully confirmed.

In order to investigate the key factors that contribute to optimal protein expression platforms, a robust reporting system is essential. Throughout the years, a variety of genetically encodable reporters have been developed to assess the efficacy of agroinfiltration. Among these, the green fluorescent protein (GFP) and its derivatives have become indispensable tools in molecular biology research (Chudakov *et al*., 2005; Lippincott-Schwartz and Patterson, 2003). However, the reliance on external light sources for visualizing fluorescence signals limits their applicability in quantitatively analyzing plants grown in natural environments. To address these limitations, alternative reporters such as the β-glucuronidase (GUS) and animal-derived luciferase enzymes have been engineered for monitoring foreign gene expression post agroinfiltration in plants (Koo *et al*., 2007). Despite their effectiveness, both the GUS and luciferase reporters necessitate the addition of costly substrates as (5-Bromo-4-chloro-3-indolyl beta-D-glucuronide) and luciferin, respectively. Moreover, the staining process for GUS or infiltration with luciferin is invasive and often results in the sacrifice of the plants or tissues under evaluation. The adoption of the pigmentation-based reporter RUBY has facilitated the noninvasive study of various plant signaling processes due to the easily observable nature of pigments (He *et al*., 2020). While this method offers convenience, it is important to note its limitations, including low sensitivity, inability for quantitation, and high background noise, which restrict its broader applicability. These drawbacks, coupled with limitations in scalability, underscore the need for further advancements in reporter systems for large-scale assessments.

Bioluminescence, the natural phenomenon of organisms emitting light, has fascinated scientists for centuries. Among many organisms that exhibit this captivating trait, bioluminescent mushrooms have garnered particular interest due to their ability to produce self-sustained visible green luminescence. This remarkable trait is made possible by the FBP, a series of enzymatic reactions that occur within the bioluminescent mushroom. Central to the FBP are four key genes including *HispS*, *H3H*, *Luz*, and *CPH*, which encode enzymes crucial for the conversion of caffeic acid, a plant metabolite, into fungal luciferin (Kotlobay *et al*., 2018). This conversion ultimately leads to the emission of light in the green spectrum, providing insight into the intricate mechanisms underlying fungal bioluminescence. This pathway has garnered attention for its potential use in creating glowing plants (Ge *et al*., 2024; Mitiouchkina *et al*., 2020; Shakhova *et al*., 2024; Zheng *et al*., 2023). Of particular interest is the self-sustained nature of caffeic acid in luminescence, making the FBP module an attractive option for plant reporting systems. Here, we identify caffeic acid as an evolutionarily conserved phenylpropanoid metabolite across land plants and introduce the FBP module as a novel reporting system. Additionally, we showcase the versatility and effectiveness of combining NahG and P19 with the FBP reporting system in effectively enhancing transient transformation in multiple plant species, and demonstrating the versatile approach for investigating gene function *in vivo*.

## Results

### FBP-based report system is applicable for all land plants

The FBP has been successfully reconstituted in plants, allowing for luminescence imaging without the need for externally supplied substrate (Calvache *et al*., 2024; Sun *et al*., 2024). This breakthrough has inspired us to develop FBP-based reporter system in plants. Given that the FBP relies on caffeic acid as a precursor for robust luminescence production, we set out to investigate whether the caffeic acid synthesis pathway is evolutionarily conserved across plant species. Our goal was to determine the feasibility of using the FBP as a reporter tool across the plant kingdom. To systematically identify potential caffeic acid synthesis pathways in plants, we analyzed caffeic acid synthesis genes from 11 high-quality genomes representing all six clades of the plant kingdom, including algae, mosses, clubmosses, ferns, gymnosperms, and angiosperms (Figure 1a and Table S1). Interestingly, our analysis revealed that all land plants possess the seven essential genes as *PAL*, *C4H*, *4CL*, *HCT*, *C3’H*, *C3H*, and *CSE*, which are responsible for converting phenylalanine to caffeic acid (Figure S1 and S2). In contrast, algae such as *Chlamydomonas reinhardtii* have lost these genes through evolution and are unable to synthesize caffeic acid (Figure 1a,b). The findings suggest that extant land plants, such as bryophytes and vascular plants, have retained the ability to produce caffeic acid as a means of adaptation throughout evolution. This phylogenetic framework further supports the potential development of FBP-based reporter systems and the creation of autoluminescent plants throughout all land plants.

**Figure 1.**
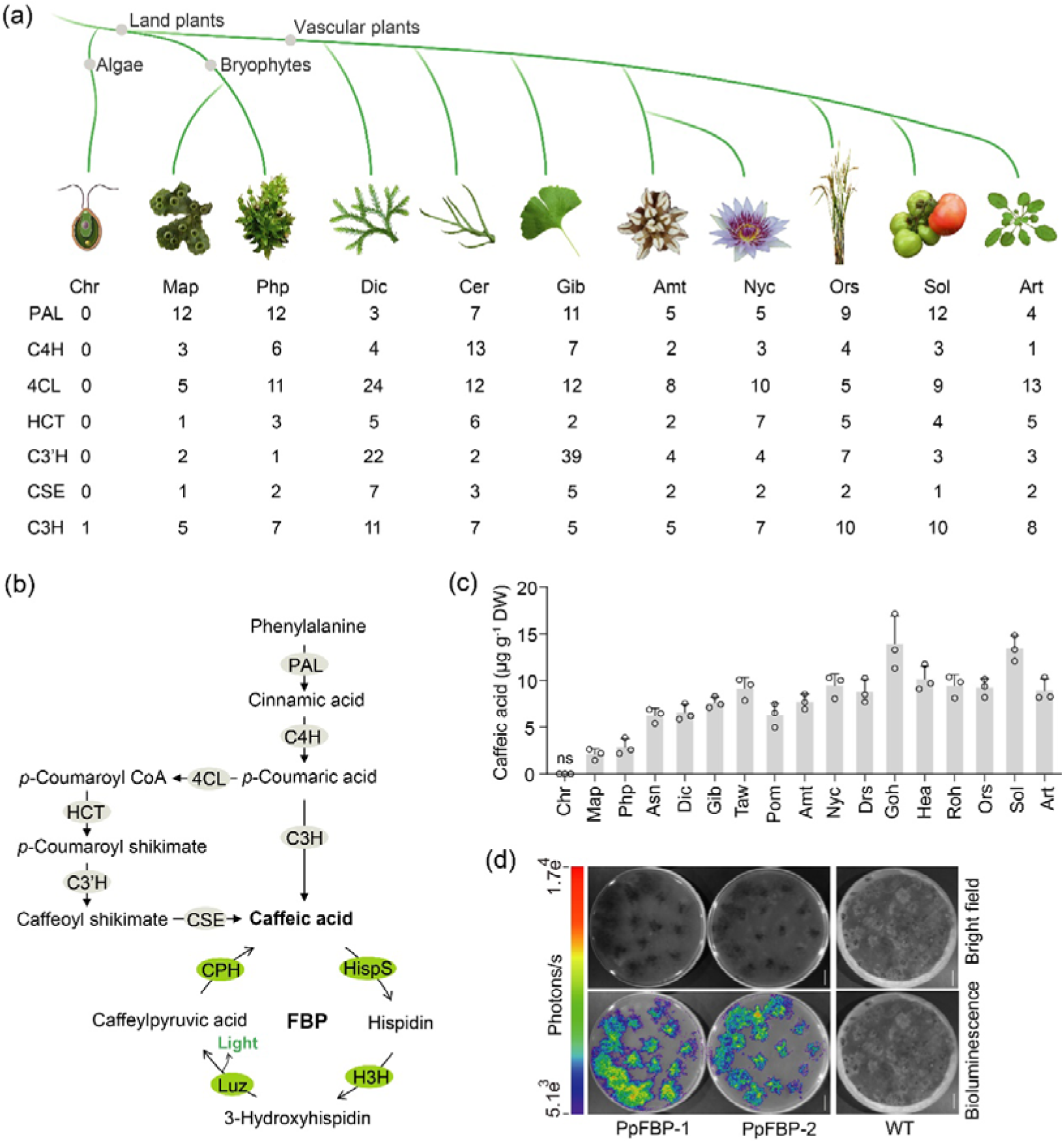
FBP precursor caffeic acid is present in all land plants. (a) Phylogenetic tree of the algae and 10 land plant species used in this study. Species marked with gray dots correspond to lateral branching. The number represents the potential number of caffeic acid synthesis related genes in each species. PAL, phenylalanine ammonia-lyase; C4H, cinnamic acid 4-hydroxylase; 4CL, 4-hydroxycinnamate: CoA ligase; HCT, 4-hydroxycinnamoyl CoA: shikimate/quinate hydroxycinnamoyl transferase; C3′H, 4-coumaroyl shikimate/quinate 3′-hydroxylase; CSE, caffeoyl shikimate esterase; C3H, coumarate 3-hydroxylase. Chr, *Chlamydomonas reinhardtii*; Map, *Marchantia polymorpha*; Php, *Physcomitrium patens*; Dic, *Diphasiastrum complanatum*; Cer, *Ceratopteris richardii*; Gib, *Ginkgo biloba*; Amt, *Amborella trichopoda*; Nyc, *Nymphaea colorata*; Ors, *Oryza sativa*; Sol, *Solanum lycopersicum*; Art, *Arabidopsis thaliana*. (b) Integrated the pathway of plant caffeic acid biosynthesis into the FBP cycle. The black arrows represent direct biochemical reactions, and FBP key genes are highlighted with the green background. (c) LC-MS/MS analysis of caffeic acid content in leaves of different species. Asn, *Asplenium nidus*; Taw, *Taxus wallichiana*; Drs, *Dracaena sanderiana*; Goh, *Gossypium hirsutum*; Hea, *Helianthus annuus*; Roh, *Rosa hybrida*. Values are mean ± SD (n = 3). ns, No signal. (d) The bioluminescent images of FBP transgenic *Physcomitrium patens*. Scale bars, 1□cm.

To assess the genetic variation in caffeic acid content among different plant species, we analyzed the levels of caffeic acid in 17 representative species and confirmed that both bryophytes and vascular plants have the ability to produce caffeic acid, while algae do not possess this compound (Figure 1c), likely due to a loss of the caffeic acid synthesis genes (Figure 1a). Interestingly, our results show that vascular plants tend to accumulate more caffeic acid compared to bryophytes (Figure 1c). In previous experiments, we successfully demonstrated the transient expression of the FBP DNA module in various plant species through *Agrobacterium* infiltration, leading to strong autoluminescence signals in *N. benthamiana*, *Phalaenopsis aphrodite*, and *Chrysanthemum morifolium*. Additionally, stable transgenic plants expressing FBP exhibited autoluminescence in *Populus canadensis* (Khakhar *et al*., 2020; Zheng *et al*., 2023), indicating that caffeic acid, derived from phenylalanine, is associated with lignin, anthocyanins, and flavonoids, which are more prevalent in vascular plants. Previous genetic and biochemical studies have suggested that although *Physcomitrella patens* does not produce lignin, homologs of the phenylpropanoid biosynthetic pathway genes are present in *Physcomitrella patens* and play a crucial role in the production of caffeate derivatives for cuticle formation (Ye and Zhong, 2022). To further explore the potential for application of the FBP system in bryophyte, we utilized Agrobacterium-mediated transformation to introduce FBP and create autoluminescent *Physcomitrella patens*. The resulting bioluminescence was observed to be weak (Figure 1d), consistent with the lower levels of caffeic acid accumulation in *Physcomitrella patens* (Figure 1c). This discrepancy underscores the genetic variations in caffeic acid content across different plant species. These results suggest that the FBP precursor caffeic acid synthesis pathway is evolutionarily conserved in plants, and the FBP-based reporter system can be effectively applied in a diverse range of land plants, particularly in vascular plants.

### Identification of enhancing factors for agroinfiltration in plants

To screen and identify the efficient enhancing factors for *Agrobacterium*-mediated transient transformation, the powerful FBP-based reporting system was utilized to identify the potential enhancing factors in agroinfiltrated leaves of *N. benthamiana*. Previous research revealed that expressing an SA hydroxylase NahG from the bacterium *Pseudomonas putida*, which breaks down SA to catechol, in *Arabidopsis* results in a notable reduction of SA. Transgenic *Arabidopsis* with overexpressed NahG displayed significantly higher GUS accumulation compared to the wild-type (Rosas-Díaz *et al*., 2017a). To investigate the impact of SA on the efficiency of agroinfiltration in *N. benthamiana*, we performed *Agrobacterium* injection with the FBP and FBP + NahG modules, respectively. Our results showed that co-overexpressing NahG resulted in higher efficiency than the control, suggesting that NahG effectively enhances the expression of foreign genes through agroinfiltration (Figure 2a). And the nuclear and cytoplasmic subcellular localization of NahG protein can be clearly observed using confocal microscopy (Figure 2b), confirming the presence of NahG in plants. Additionally, we assessed whether the enhanced efficiency of *Agrobacterium*-mediated transient expression with the *NahG* gene is dependent on the reduced accumulation of SA. Our measurements indicated that leaves agroinfiltrated with the NahG exhibited significantly lower levels of SA (Figure 2c), supporting the notion that the improved transformation with NahG is correlated with decreased SA content. Given the role of SA as a crucial signaling molecule in plant immunity and its ability to confer resistance against various pathogens, strategies such as overexpressing NahG that can circumvent and suppress immune responses have the potential to greatly enhance the transient expression of foreign genes.

**Figure 2.**
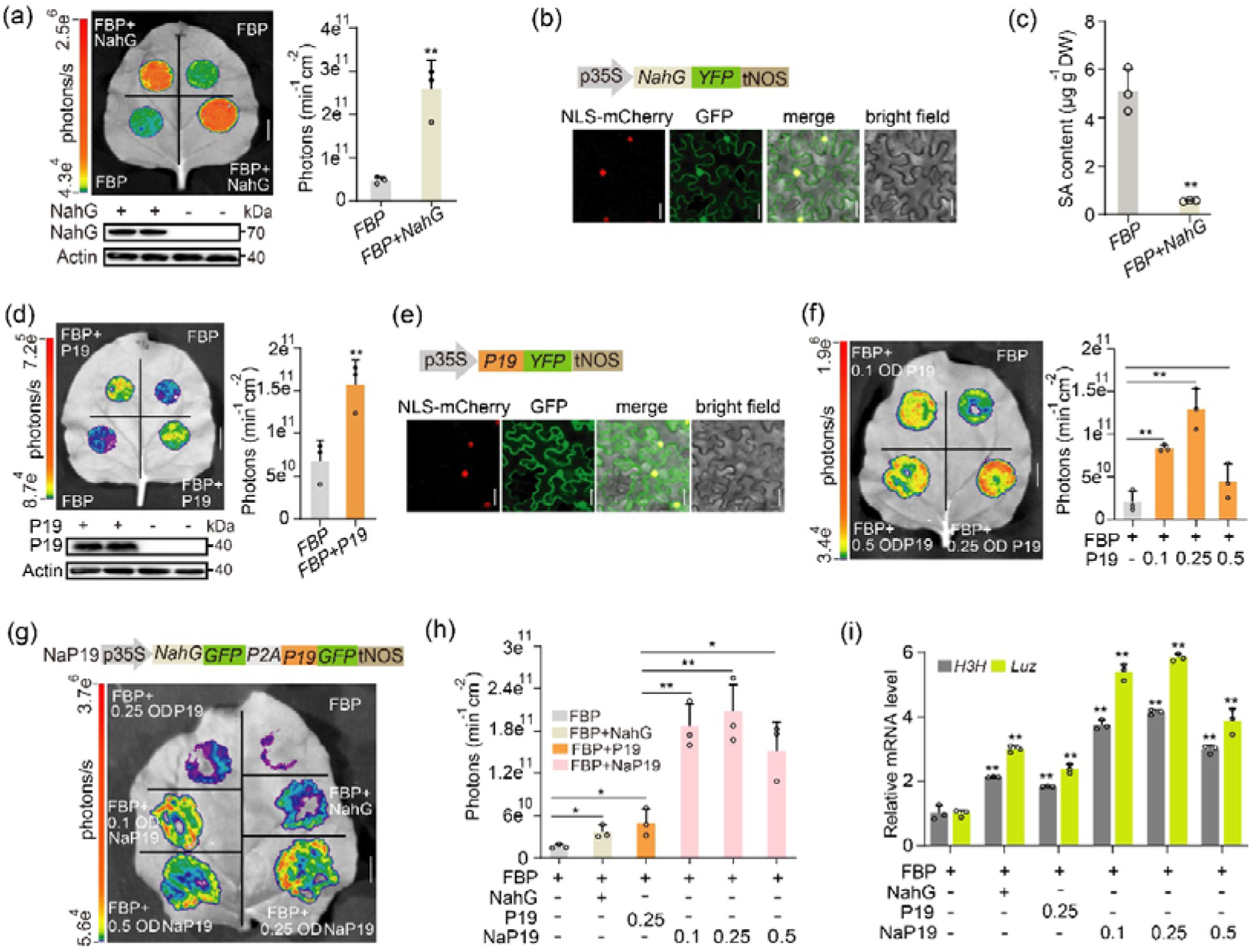
Identification of *NahG* and *P19* effectively enhances agroinfiltration efficiency with the FBP report system. (a) NahG intensifies the photons emitted from FBP-infiltrated *N. benthamiana* leaves. Bioluminescent images were collected 72 h after *Agrobacterium* infection. NahG transiently expressed in tobacco leaves was determined by immunoblot analyses. Values are mean ± SD (n = 3). Statistical significance was assessed using two-tailed Student’s *t*-tests (\*\**P* ≤ 0.01). (b) Subcellular localization of NahG, NLS-mCherry was used as nuclear marker. Scale bars, 20 μm. (c) LC-MS/MS analysis of the content of Salicylic acid (SA) from injected tobacco leaves. Values are mean ± SD (n = 3). Statistical significance was assessed using two-tailed Student’s *t*-tests (\*\**P* ≤ 0.01). (d) P19 enhances the expression of FBP in infiltrated *N. benthamiana* leaves, bioluminescent intensity and protein expression analyses were carried out 72 h after *Agrobacterium* infection. Western blot analysis was conducted using GFP antibody. Values are mean ± SD (n = 3). Statistical significance was assessed using two-tailed Student’s *t*-tests (\*\**P* ≤ 0.01). (e) Subcellular localization of P19, NLS-mCherry was used as nuclear marker. Scale bars, 20 μm. (f) Analysis of agroinfiltration enhancement of P19 at different concentrations, bioluminescent intensity analysis was conducted 72 h after *Agrobacterium* infection. The OD_600_ values of P19 shown above are labeled as the final concentration of the infection solution. Values are mean ± SD (n = 3). Statistical significance was assessed using two-tailed Student’s *t*-tests (\*\**P* ≤ 0.01). (g-i) Analysis to determine the enhancement effect of NahG and various concentrations of P19 in agroinfiltration. Bioluminescent imaging (g), photon analysis (h), and qPCR assays (i) on H3H and Luz were performed 72 h after infection with Agrobacterium. NaP19 indicating the NahG-P2A-P19. Scale bars, 1□cm. Values are mean ± SD (n = 3). Statistical significance was assessed using two-tailed Student’s *t*-tests (\*\**P* ≤ 0.01).

Several *P. syringae* type III effectors, such as AvrPto and AvrPtoB, have been shown to suppress the plant’s basal defense system and block multiple PAMP-mediated signaling pathways (Oh and Martin, 2011). Additionally, other effectors from the *P. syringae* family, like HopF2 and AvrPphE, inhibit host immunity by breaking down JAZs and BAK signaling proteins (Sarris *et al*., 2011). These effectors potentially enhance the expression of foreign genes in T-DNA by hindering the plant’s natural defense mechanisms. To evaluate these effectors, we compared them with NahG in agroinfiltrated leaves of *N. benthamiana*, and observed a significant decrease in normalized bioluminescence signals in the presence of type III effectors (Figure S3a-d), indicating that they do not transiently boost *Agrobacterium*-mediated infection in plants, unlike stable transformation methods. Furthermore, the chromatin remodeling effector GIF1 did not significantly enhance transient transformation efficiency compared to NahG, as indicated by the levels of bioluminescence produced by FBP (Figure S3a,b). We attempted to enhance transient transformation efficiency by co-expressing HopF2 and AvrPtoB in *N. benthamiana*, but no significant difference was detected (Figure S3c,d). In contrast, delivery of NahG increased agroinfiltrated efficiency by more than two-fold (Figure S3c,d). This trend was validated by the successful delivery of P. syringae type III effectors and GIF1 into *N. tabacum* and *Gossypium barbadense*, respectively. Interestingly, only NahG significantly enhanced *Agrobacterium*-mediated transient infection in multiple plant species, as shown in Figure S3e,f. This observation revealed a distinct difference in the mechanisms involved in agroinfiltration and stable transformation in plants. The findings also underscore the importance of understanding the specific roles of enhancing factors in these processes.

Previous studies have demonstrated that the co-expressed silencing inhibitor P19 from tomato bushy stunt virus can suppress PTGS by sequestering siRNAs, thereby enhancing the expression of *GFP* (Saxena *et al*., 2011). Here, we aimed to evaluate the efficacy of P19 in enhancing the transient expression of foreign genes in plants. To accomplish this, we employed an FBP reporter in an agroinfiltration assay, co-expressing FBP and P19 in *N. benthamiana* leaves. Our results revealed a significant two-fold increase in bioluminescence signal in the presence of P19 (Figure 2d). Additionally, we investigated the subcellular localization of P19 in heterologous hosts using confocal microscopy, confirming nuclear and cytoplasmic distribution (Figure 2e). Considering the impact of P19 expression levels on plant developmental processes (Hamel *et al*., 2024), we conducted a concentration analysis and determined that an OD_600_ of 0.25 of *Agrobacterium* carrying P19 effectively improved the transient expression efficiency of foreign genes (Figure 2f). Our study suggests that P19, when expressed at the appropriate level, has the potential to enhance the transient expression of foreign genes in plants by reducing PTGS. This indicates that P19 could be valuable in a variety of plant biotechnological applications, including genome editing, transgenic technology, and synthetic biology.

The combined effects of co-expressing different enhancers on suppressing the host plant’s immune response and PTGS to enhance foreign genes expression in agroinfiltrated *N. benthamiana* have not been investigated before. In this resource study, we designed a co-expression construct, NaP19, by linking NahG and P19 using the self-cleaving peptide sequence P2A. Through Agrobacterium-mediated transient expression assays, we observed a significant tenfold increase in photo flux when NaP19 (NahG+P19) was co-overexpressed at an OD_600_ concentration of 0.25 compared to FBP alone (Figure 2g,h). These data also indicated that NaP19 dramatically improved transient expression efficiency compared to expressing NahG or P19 individually (Figure 2g,h). Since FBP intensity can be influenced by endogenous metabolic processes and external environment factors (Ge *et al*., 2024; Mitiouchkina *et al*., 2020), subsequent qPCR analysis confirmed that the enhanced photo intensity was due to increased expression of foreign genes *H3H* and *Luz* in the FBP reporter system (Figure 2i). These results further suggest that the FBP reporter system is highly effective, as demonstrated by the positive correlation between bioluminescence and transient expression efficiency. Overall, our findings support the notion that NahG and P19 may synergistically enhance agroinfiltration efficiency.

### NaP19 facilitated agroinfiltration in multiple species

To investigate the effectiveness of NaP19-mediated agroinfiltration in various plant species, we conducted experiments using nine different plant species, including ornamental plants such as *Adenium obesum*, *Dahlia pinnata*, *Tulipa gesneriana*, *Begonia hiemalis*, *Rhamnus hybrida*, *Phalaenopsis aphrodite*, as well as main crops like *Helianthus annuus*, *Gossypium hirsutum* and *Vigna radiate*. Despite the importance of these plants in horticulture and agriculture, the use of *Agrobacterium*-mediated transient expression assays has not been well-established, limiting molecular research and genetic improvement efforts in these plants. In this study, we performed agroinfiltration experiments using *Agrobacterium* cultures carrying the FBP and NaP19 constructs, respectively. Bioluminescence signals were detected following agroinfiltration of multiple plant species, and the expression levels of *H3H* and *Luz* transcripts were quantified using qPCR at the third post-inoculation. Our analysis demonstrated that plants infiltrated with NaP19 exhibited a significant increase in FBP expression and emitted more intense bioluminescence signals across various plant species, as compared to plants that were solely agroinfiltrated with FBP and empty vectors (Figure 3a-h and Figure S5a-i). Subsequent transcript assays further confirmed elevated levels of FBP transcripts in multiple plant tissues with the NaP19-mediated approach (Figure 3a-i). The correlation between the accumulation of FBP transcripts and bioluminescence signals further supports the efficiency of the FBP reporter system in vascular plants. Additionally, our findings suggest that NaP19 can facilitate *Agrobacterium*-mediated transient expression in a wide range of plant species.

**Figure 3.**
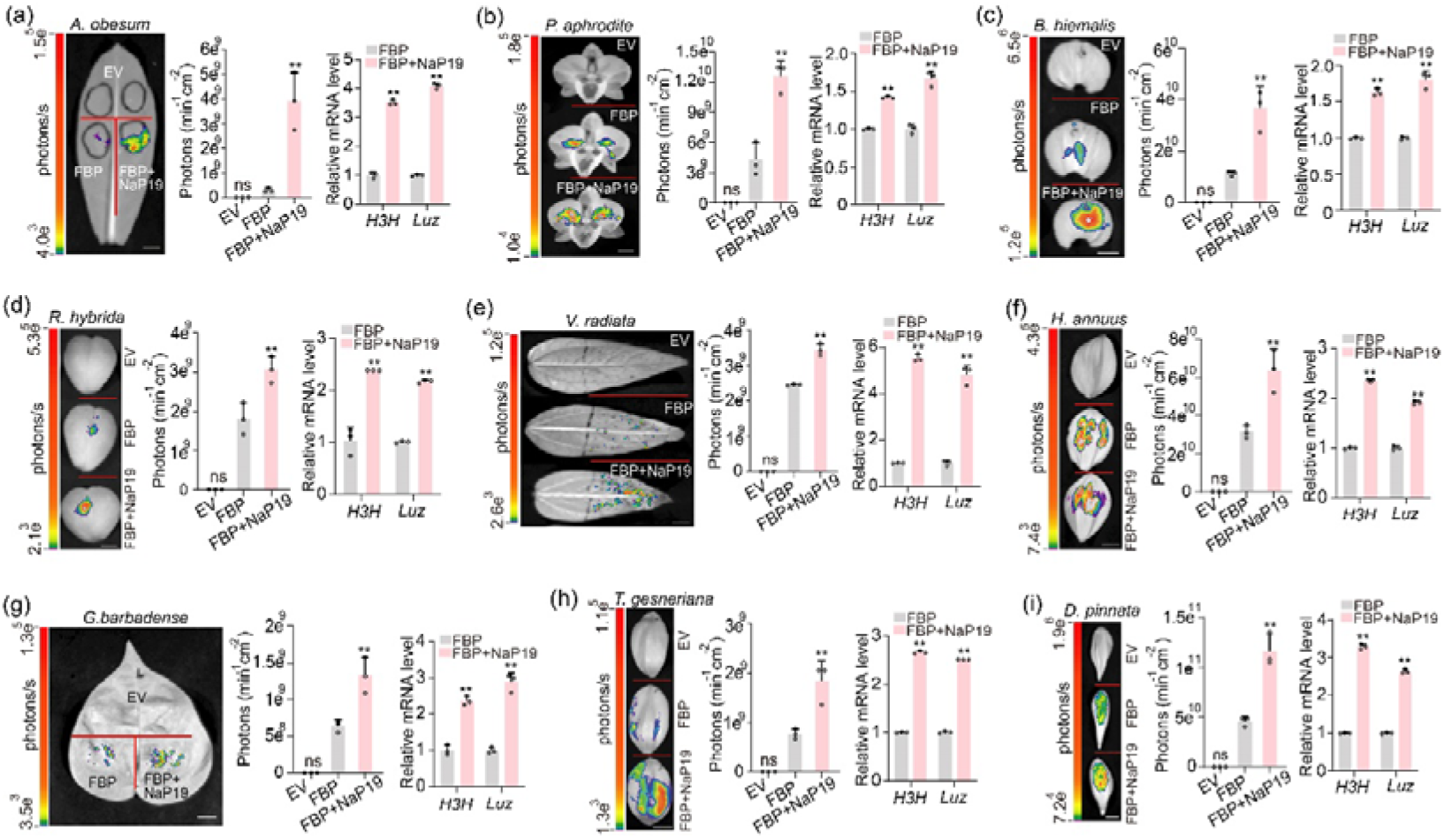
The expression of NaP19 enhances the efficiency of agroinfiltration in various plant species with the FBP reporter system. (a-h) The bioluminescent intensity analysis and RT-qPCR assay were conducted on infiltrated tissues of various plant species, including *Adenium obesum* (a), *Phalaenopsis aphrodite T* (b), *Begonia hiemalis* (c), *Rhamnus hybrida* (d), *Vigna radiate* (e), *Helianthus annuus* (f), *Gossypium hirsutum* (g), *Tulipa gesneriana* (h), and *Dahlia pinnata* (i). The tissues were treated with different components, including control (CT), FBP, and FBP + NaP19 after 72 h. Reference genes for each species can be found in Table S2. Scale bars, 1□cm. ns, No signal. Values are mean ± SD (n = 3). Statistical significance was assessed using two-tailed Student’s *t*-tests (\*\**P* ≤ 0.01).

Through our research, we found that the NaP19 genes had a positive impact on transient gene delivery systems, as evidenced by improvements at both the metabolic and mRNA levels (Figure 3 and Figure S5). Building on these promising results, we expanded our analysis to a diverse range of vascular plants including *Syzygium grijsii*, *Senecio radicans*, *Ranunculus asiaticus*, *Consolida ajacis*, *Bryophyllum pinnatum*, *Cyclamen persicum*, *Dracaena sanderiana*, *Campanula medium*, *Ligustrum ovalifolium*. By co-infiltrating plant tissues with *Agrobacterium* cultures containing both the FBP and NaP19 expression vectors, respectively, we observed a stronger bioluminescence signal compared to samples infiltrated with the FBP alone (Figure 4a-i). Furthermore, we broadened our investigation to include gymnosperms like *Nageia nagi*, and *Taxus chinensis*, as well as angiosperms encompassing dicots (*Rosa odorata*, *Camellia japonica*, *Freesia hybrida*, *Prunus serrulata*, *Epipremnum aureum*, *Tagetes erecta*, *Catharanthus roseus*, *Taxus chinensis*, *Michelia figo*, *Nageia nagi*) and monocot *Freesia hybrid* (Figure S5a-j). We observed a significant increase in bioluminescence in plants co-infiltrated with *Agrobacterium* cultures carrying both the FBP and NaP19 expression vectors. The findings demonstrate that our approach effectively enhances the agroinfiltration technology in a wide variety of plant species within the vascular plants group. This highlights the versatility of the FBP reporter system in all vascular plants and emphasizes the crucial role of NaP19 in establishing an efficient agroinfiltration approach across most plant tissues *in vivo*, thus benefiting plant science and molecular pharming across various plant species.

**Figure 4.**
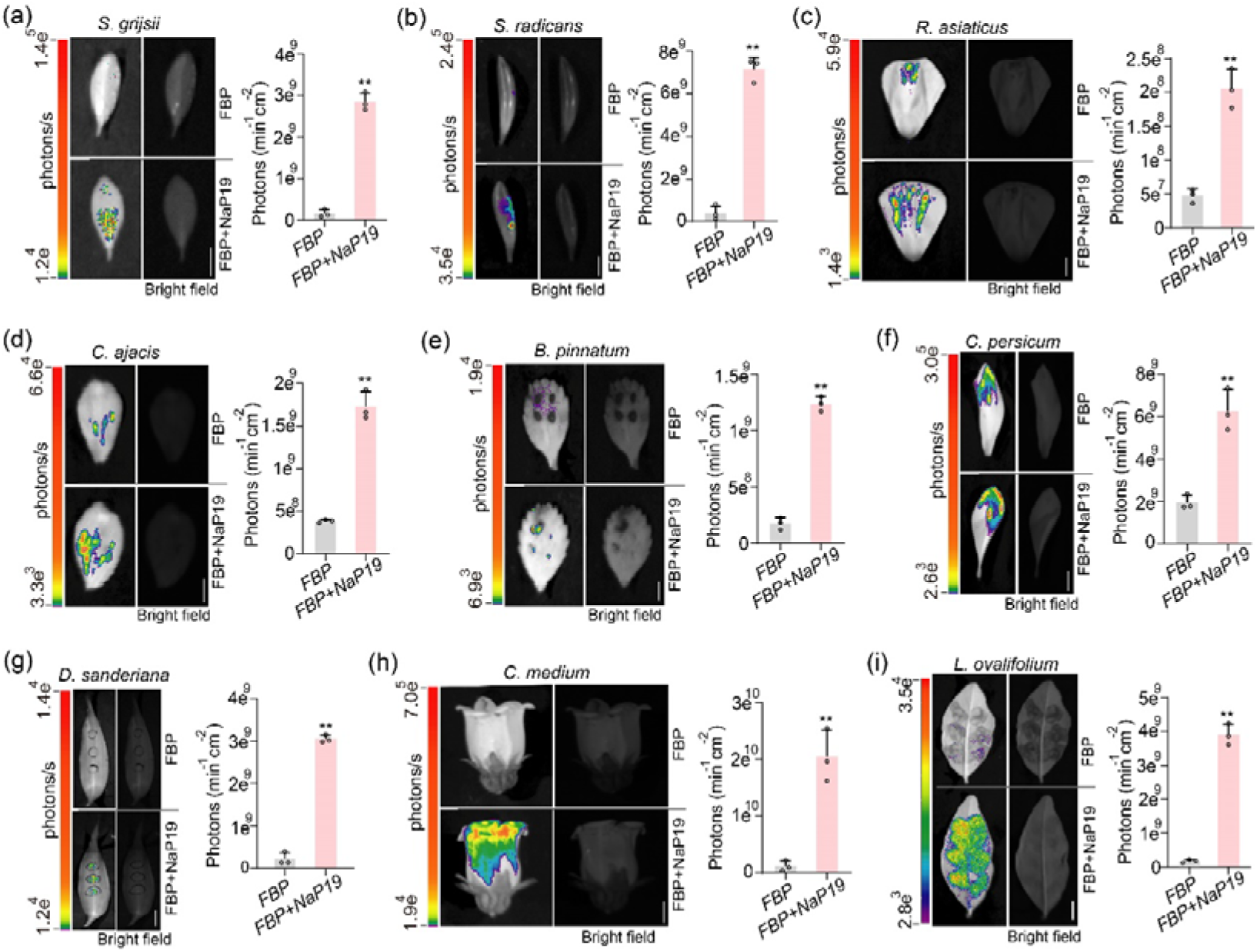
The test of NaP19 enhances the agroinfiltration efficiency with the *FBP* report system in diverse plant species. (a-i) Agroinfiltration of *Syzygium grijsii* (a), *Senecio radicans* (b), *Ranunculus asiaticus* (c), *Consolida ajacis* (d), *Bryophyllum pinnatum* (e), *Cyclamen persicum* (f), *Dracaena sanderiana* (g), *Campanula medium* (h), *Ligustrum ovalifolium* (i) tissues with FBP and FBP+NaP19 modules respectively. Bioluminescent intensity analysis from agroinfiltrated leaves after 72□h. Scale bars, 1□cm. Error bars indicate means ± SD (n□=□3). Statistical significance was assessed using two-tailed Student’s *t*-tests (\**P*□≤□0.05, \*\**P*□≤□0.01).

### Using the agroinfiltration method for *in vivo* assay of sunflower and mung bean

Expanding on our above successful co-infiltration experiments with FBP and NaP19 in *Helianthus annuus* and *Vigna radiata* tissues (Figure 3e,f), we further aimed to develop an agroinfiltration approach specifically for *in vivo* bioassays in *Helianthus annuus*. By introducing nuclear localization sequence (NLS)-mCherry *Helianthus annuus* tissues using NaP19-mediated agroinfiltration (Figure 5a), we sought to assess the adaptability of the approach for investigating protein subcellular localization in *Helianthus annuus*. Confocal microscopy results depicted a robust nuclear signal in *Helianthus annuus* (Figure 5b), indicating a successful transformation with our novel approach. Additionally, western blot analysis of NahG protein further confirmed the efficient expression of NaP19 in Helianthus annuus (Figure 5c). These findings collectively support the efficacy of the NaP19-mediated agroinfiltration method in *Helianthus annuus* tissues for *in vivo* assays.

**Figure 5.**
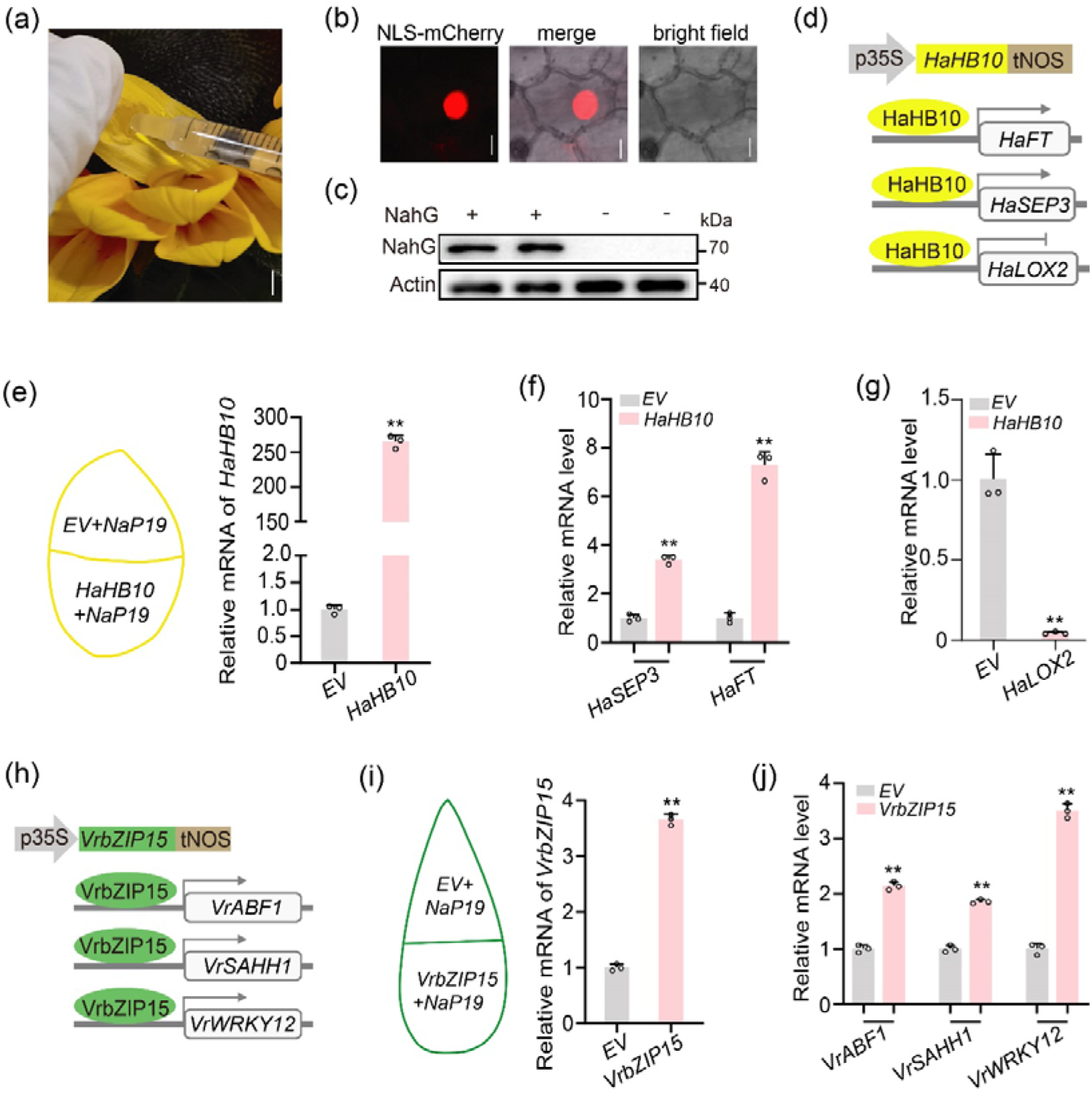
Optimized agroinfiltrated strategies for investigating transcriptional regulation in sunflower and mung beans. (a) Schematic diagram of sunflower petal injection using the syringe. Samples from flat petals of sunflower. Scale bar, 1□cm. (b) Subcellular localization of NLS-mCherry visualized using a Zeiss LSM 880 confocal microscope. Scale bars, 10 μm. (c) NahG from NaP19 transiently expressed in tobacco leaves was determined by immunoblot analyses using GFP antibodies. (d) Schematic diagram of the *HaHB10* expression constructs. HaHB10 acts upstream of *HaSEP3*, *HaFT* and *HaLOX2*. (e) qPCR assay of HaHB10 in agroinfiltrated sunflower petals, empty vector (EV) as control illustrated in the diagram. Values are mean ± SD (n = 3). Statistical significance was assessed using two-tailed Student’s *t*-tests (\*\**P* ≤ 0.01). (f and g) qPCR assay of upregulated genes *HaSEP3*, *HaFT* (f) and downregulated gene *HaLOX2* (g). Values are mean ± SD (n = 3). Statistical significance was assessed using two-tailed Student’s *t*-tests (\*\**P* ≤ 0.01). (h) A schematic diagram illustrating the regulatory relationship between VrbZIP15 and VrABF1, VrSAHH1, and VrWRKY12 *in vivo*. (i and j) qPCR assay of the overexpression of *VrbZIP15* (i) and the upregulation genes *VrABF1*, *VrSAHH1*, and *VrWRKY12* (j) *in vivo*. Values are mean ± SD (n = 3). Statistical significance was assessed using two-tailed Student’s *t*-tests (\*\**P* ≤ 0.01).

Transcriptional regulation experiments are a commonly used method to build interaction networks that offer valuable insights into transcription factors, target genes, and biological processes in organisms. In this study, we focused on the transcription factor HaHB10, an HD-Zip II transcription factor in sunflowers known for its role in flowering induction and the regulation of SA and jasmonic acid responses (Dezar *et al*., 2011). Previous studies on *Helianthus annuus* genes have typically relied on stable transformation assays in *Arabidopsis* due to the lack of an efficient stable transformation system in *Helianthus annuus*. To overcome this limitation, we adopted the agroinfiltration method to co-express *HaHB10* and *NaP19* constructs (Figure 5d) and performed qPCR analysis to validate the overexpression of HaHB10 in *Helianthus annuus* tissues (Figure 5e). Through this innovative approach, we successfully identified downstream genes regulated by HaHB10. Notably, we observed a significant upregulation of the flowering-related genes E-class *SEPALLATA* (*SEP*) and *FLOWERING LOCUS T* (*FT*) in *Helianthus annuus* tissues upon HaHB10 overexpression (Figure 5f). Conversely, the gene *HaLOX2*, involved in jasmonic acid biosynthesis, was downregulated by HaHB10 (Figure 5g). Our *in vivo* molecular analyses demonstrated that the NaP19-assisted agroinfiltration method is a powerful tool for studying protein subcellular localization and transcriptional regulation. This method allows for the investigation of the impact of regulatory genes on plant transcriptional networks without the need for stable transformation processes, making it particularly valuable for plant species lacking stable transformation technology.

The effectiveness of agroinfiltration technology in *Vigna radiata* for molecular biology analysis is yet to be determined. Here, we sought to evaluate the potential of the NaP19-mediated agroinfiltration method in *Vigna radiata* tissues by introducing NaP19 and unidentified bZIP transcription factor VrbZIP15 (Figure 5h), a homolog of *Glycine max* GmbZIP15 known for its involvement in salinity and drought stress response (Zhang *et al*., 2020a). Through qPCR analysis, we confirmed the successful expression of *VrbZIP15* in *Vigna radiata* (Figure 5i) and investigated the expression levels of several potential downregulated genes by predication based on the homology analysis in *Glycine max*. The predicted genes as *VrABF1*, *VrSAHH1*, and *VrWRKY12*, which encode a bZIP transcription factor, phosphate dehydrogenase, and WRKY transcription factor, respectively. Our results revealed a significant upregulation in the expression of these three genes (Figure 5j), indicating a potential transcriptional regulation relationship with the VrbZIP15 pathway in *Vigna radiata*. Overall, our findings demonstrate the applicability of the NaP19-mediated agroinfiltration technique for examining protein subcellular localization and transcriptional regulation in crops like *Helianthus annuus* and *Vigna radiata*. Notably, the technology eliminates the need for lengthy stable transformation procedures, setting the stage for further research in molecular biology and synthetic biology in multiple species.

### NaP19 mediated agroinfiltration *in vivo* assay in multiple cotton cultivated species

Our study aimed to evaluate the efficacy of an optimized agroinfiltration method in cotton, a crucial crop globally and particularly in China, where traditional methods have proven challenging to apply across different genotypes. In our investigation, we found that the use of NaP19 significantly improved the efficiency of agroinfiltration in cotton, as illustrated in Figure 3g. Specifically, we tested this method on two cultivated allotetraploid *Gossypium* species including cultivars of *Gossypium hirsutum,* acc. Jin668, *Gossypium barbadense,* acc. Hai7124, and 3-79 to assess their susceptibility to agroinfection and expression of foreign genes. Our results, depicted in Figure 6a, show that the GhbHLH121-GFP fusion protein produced consistent GFP signals in the nucleus and cytoplasm of all three cotton genotypes, mirroring results seen in *N. benthamiana* (Li *et al*., 2023). Moreover, using NaP19-mediated agroinfiltration, we delved into the interaction between iron regulatory transcription factor bHLH121 and POPEYE (PYE)/bHLH47 in response to iron deficiency in the *Gossypium* species. By employing their respective homologues in cotton GhbHLH121 and GhPYE (Gao *et al*., 2020), we successfully demonstrated protein-protein interactions (PPI) through a luciferase complementation assay with NaP19 assistance (Figure 6b). Our findings confirmed the interaction between GhbHLH121 and PYE in cotton leaves, as evidenced by the restoration of bioluminescence in case of constructs expressing PYE-nLuc and cLuc-bHLH121. Overall, our study highlights the efficacy and versatility of the NaP19-mediated agroinfiltration method for protein localization and interaction assays in diverse cotton varieties, underscoring its potential to overcome genotype limitations in various plant species.

**Figure 6.**
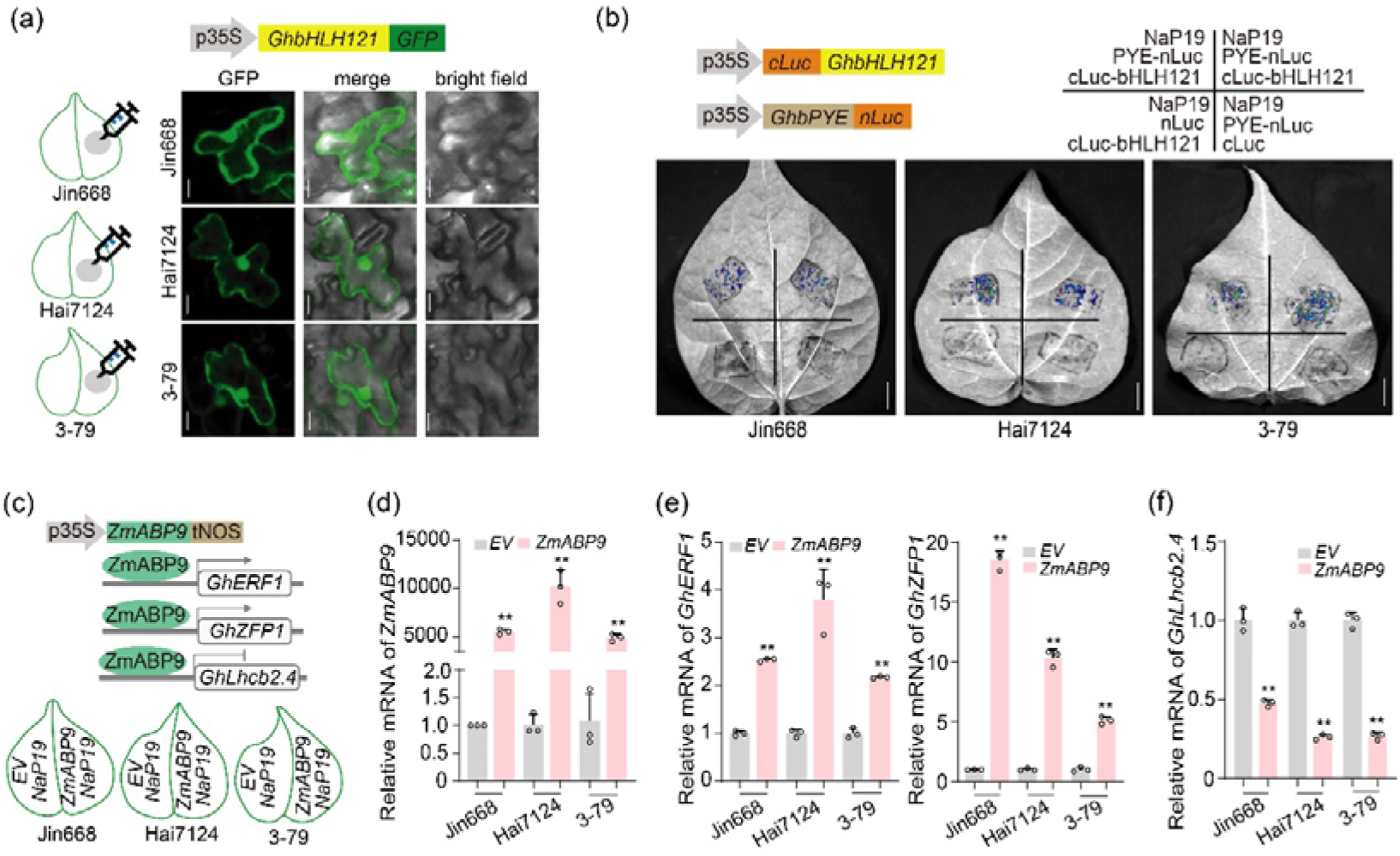
NaP19-mediated transient transformation method for assessing luciferase complementation and transcriptional regulation assay in various cotton species. (a) The subcellular localization of GhbHLH121 in *Gossypium hirsutum*, acc Jin668 and *Gossypium barbadense*, acc Hai7124 and 3-79 were visualized using a confocal microscope. Scale bars, 10 μm. (b) The Luciferase complementation assay of the interactions between GhbHLH121 and GhPYE. Fusion vectors were introduced into cotton leaves for transient coexpression. Scale bar, 1□cm. (c) The schematic illustrations of ZmABP9 stimulate the expression of *GhERF1*, *GhZEP1* and *GhLhcb2.4*. The injection diagram indicates the agroinfiltration of devious cotton species. EV as control. (d-f) qPCR assay of the expression of *ZmABP9* (d) upregulated genes *GhERF1* and *GhZEP1* (e), downregulated gene *GhLhcb2.4* (f) in agroinfiltrated cotton leaves. Values are mean ± SD (n = 3). Statistical significance was assessed using two-tailed Student’s *t*-tests (\*\**P* ≤ 0.01).

We further explored the transcriptional regulation network in different cotton genotypes by introducing the maize gene ZmABP9, known for its stress tolerance enhancement and involvement in the ABA signaling pathway (Zhang *et al*., 2011). Previous studies have shown that ZmABP9 can increase the expression of stress-related genes such as *GhZFP1*, *GhERF1*, and *GhLhcb2.4* under salt stress in stably transformed cotton lines overexpressing ZmABP9 (Wang *et al*., 2017). To streamline the analysis, we utilized a transient assay through NaP19-mediated agroinfiltration technology (Figure 6c) and performed qPCR to validate the overexpression of ZmABP9 in three distinct cotton genotypes (Figure 6d). Subsequent qPCR assessments unveiled the upregulation of *GhZFP1* and *GhERF1* by ZmABP9 in all three cotton accessions (Figure 6e), alongside the noticeable downregulation of *GhLhcb2.4* (Figure 6f). These results suggest that the NaP19-mediated agroinfiltration approach exhibits sufficient efficiency and sensitivity in molecular-level investigations across diverse cotton accessions. Overall, our findings underscore the efficacy of the NaP19-mediated agroinfiltration method in diverse cotton genotypes, regardless of their genetic backgrounds. This innovative approach shows great potential in advancing gene characterization and synthetic biology applications in various plant species.

## Discussion

Agroinfiltration is a valuable method for transient gene expression in plants, particularly in *N. benthamiana*. However, its extensive application is limited to this plant species, which is commonly used in synthetic and molecular biology research. To expand its utility to a broader range of plants, it is necessary to optimize its capabilities. One major challenge is the absence of a universal reporter system that can reliably assess efficiency across different plant species. Previous studies have relied on GUS as a reporter, but its sensitivity can vary based on the substrate used, leading to inconsistent results (Cervera, 2005; Jefferson, 1989). Additionally, GUS and fluorescent protein reporters have drawbacks in sensitivity and quantitative analysis (Baulcombe *et al*., 1995; Cui *et al*., 2016). The recently introduced RUBY system, comprising three genes, has shown promise as an effective, non-invasive, and cost-effective tool for monitoring gene expression and plant transformation (He *et al*., 2020; Wang *et al*., 2023). Despite their advantages, RUBY-based assays are limited by sensitivity, quantification issues, and high background levels in various plant species.

The discovery of the FBP from glowing fungus presents new opportunities for enhancing reporter systems in biological assays (Calvache *et al*., 2024; Kotlobay *et al*., 2018). The FBP, involving four enzymes (HispS, H3H, Luz, and CPH), has a cyclic metabolic route (Figure 1b). The synthesis pathway of caffeic acid has evolutionarily conserved homologs in all land plants (Figure 1, and Figure S1,2), suggesting that plant cells may utilize their own caffeic acid pool to support the FBP independently, without requiring additional substrate. This discovery highlights the potential of the FBP as a valuable reporter in plants for visualizing transient transformation events more conveniently. In this study we have successfully introduced the FBP into multiple plant species, enabling accurate and quantitative bioluminescence detection using an ultra-sensitive charge coupled device (CCD) camera, without additional substrate or excitation light sources. Results (Figure 3,4, and Figure S5) demonstrate the advantages of the FBP system over traditional GUS or GFP reporter systems. The FBP was effective in all land plants and facilitated the screening of transient transformation events with varying enhancement factors, accurately assessing efficiency and avoiding false negative results. Leveraging the FBP reporter holds promise for enhancing molecular biology and molecular pharming in plant species.

Agroinfiltrated tissues in host plants trigger a basal immune response similar to pattern-triggered immunity by microbe-associated molecular patterns (Zhang and Zhou, 2010). This immune response obviously impedes *Agrobacterium*-mediated transformation efficiency (Pitzschke, 2013). To address this issue, strategies can be implemented to prevent or suppress immune reactions in *N. benthamiana* plants post-agroinfiltration. One approach involves using the bacterial type-III effector AvrPto to inhibit immune-related kinases. Studies have shown that modifying *Agrobacterium* by introducing *P. syringae* type III effectors AvrPto, AvrPtoB, or HopAO1 can enhance stable transformation efficiencies in tobacco, wheat, and alfalfa (Raman *et al*., 2022). However, certain *P. syringae* effectors like AvrPto, AvrPtoB, HopF2, AvrPphE, and chromatin remodeling factor GIF1 do not improve agroinfiltration efficiency (Figure S3), indicating differences between stable transformation and agroinfiltration processes. SA is a key defense phytohormone, plays a crucial role in basal defense mechanisms, amplifying local immune responses and establishing systemic acquired resistance (Ngou *et al*., 2022; Spoel and Dong, 2024). In our study, overexpression NahG significantly reduced SA levels in plants, leading to improved transient expression efficiencies (Figure 2a-c) by preventing or suppressing host plants immune responses. Agroinfiltration often triggers a PTGS response, leading to the destruction of transgene-encoded mRNA (Hamilton *et al*., 2002; Voinnet *et al*., 2000). Strategies such as the removal of RNA-dependent RNA polymerase 6 and co-expression of the silencing inhibitor P19 have been employed to counteract PTGS and enhance transgene expression (Matsuo and Atsumi, 2019). Our study found that utilizing the P19 significantly increased bioluminescence and expression of *Luz* and *H3H* in FBP (Figure 2d-f). The optimal concentration of P19 for maximum FBP yield was determined to be 0.25 OD (Figure 2F and 2G), with excessive stable expression leading to leaf tissue developmental defects (Mao *et al*., 2018), underscoring the importance of optimizing the concentration and duration of P19 treatment. By further combining P19 with NahG during agroinfiltration showed significantly improved efficiency in FBP expression compared to using them individually (Figure 2g-i and Figure 3). The data conclusively demonstrate the efficacy of an integrative approach combining strategies to minimize immune responses and RNA degradation for optimizing agroinfiltration with extensive application.

Transient expression technologies such as agroinfiltration and plant protoplast transformation have been utilized for rapid gene testing for decades (Jay *et al*., 2023). However, protoplast methods are complicated and rely on the proficiency of the experimentalist to generate high-quality protoplasts, with concerns about cell viability post cell wall removal for *in vivo* assays (Reyna-Llorens *et al*., 2023; Yoo *et al*., 2007). Although agroinfiltration is beneficial, its effectiveness is often hindered by species compatibility issues, especially in non-*N. benthamiana* species (Zhang *et al*., 2020b). To address these limitations, a new transient transformation method utilizing NaP19 has been created. This innovative approach significantly improves transformation efficiency across a wide range of plant species, (Figure 3,4 and Figure S4,5), including those not previously amenable to agroinfiltration technologies such as horticultural plants and perennial trees. This method has been successfully applied to study transcriptional regulation processes and PPI in cash crops like *Helianthus annuus*, *Vigna radiata*, and various cotton varieties (Figure 5,6), overcoming genotype-specific constraints. Furthermore, this technique will allow for the exploration of genetic circuits and metabolic engineering in diverse plant species. By integrating CRISPR activation and interference technologies, there is potential to utilize host plants for producing plant-specific secondary metabolites in other species, such as paclitaxel and vinblastine. In conclusion, our optimized agroinfiltration systems offer significant advantages over stable transformation methods and protoplast assays for a variety of research applications. The efficacy of the FBP reporter system in advancing plant biotechnology in all land plants underscores the numerous benefits of this innovative reporter. Our study not only emphasizes the wide-ranging advantages of this technology, but also highlights the potential for exploiting agroinfiltration technologies in plant synthetic biology.

## METHOD DETAILS

### Plasmid construction

In this study, we generated the plasmids used by sequentially cloning various genetic elements into the pCAMBIA1300 vector. First, the 35S promoter (p35S) and NOS terminator (tNOS) sequences were inserted into the pCAMBIA1300 plasmid using Hind III/Xba I and Sac I/EcoR I restriction enzymes, respectively. Next, the *YFP* and *eGFP* fragments were added to the pCAMBIA1300-p35S vector using BamH I/Sac I enzymes, resulting in the generation of pCAMBIA1300-p35S-YFP and pCAMBIA1300-p35S-eGFP plasmids. The *P19* gene (GenBank ID: AJ288942.1) from tomato bushy stunt virus and the *NahG* gene (GenBank ID: M60055) from *Pseudomonas putida* were then amplified and connected using Overlap Extension PCR with a 2A peptide linker. Subsequently, the amplified *P19* and *NahG* fragments were seamlessly cloned into the pCAMBIA1300-p35S vector using 2X MultiF Seamless reagent (ABclonal, USA), resulting in the formation of pCAMBIA1300-p35S-NahG-YFP, pCAMBIA1300-p35S-P19-YFP, and pCAMBIA1300-p35S-NahG-2A-P19 plasmids. Similarly, the pCAMBIA1300-p35S-GhbHLH121 and pCAMBIA1300-p35S-VrbZIP15 constructs were generated using the same cloning methodology. To construct vectors that enhance infection, fragments such as p35S-Avrpto-tNOS and p35S-AvrptoB-tNOS were inserted into the pCAMBIA1300-NahG-YFP using the HindIII restriction enzyme. Similar procedures were followed for the pCAMBIA1300-HopF2-NahG, pCAMBIA1300-AvrPphe-NahG, and pCAMBIA1300-GIF1-NahG vectors. In the case of the three factor vectors, the HopF2 fragment was inserted into the pCAMBIA1300-AvrptoB-NahG using the KpnI restriction enzyme.

The FBP constructs contained the codon-optimized *NnLuz*, *NnHispS*, *NnH3H*, and *NnCPH* genes from *N. nambi*, as well as the *AnNPGA* gene from *A. nidulans*. The assembly of the FBP constructs was performed using the TransGene Stacking II system, as detailed in previous reports (Zheng *et al*., 2023; Zhu *et al*., 2017). The primer sequences utilized in the plasmid constructions can be found in Table S2, the vectors information is shown in Table S3.

#### Agrobacterium-mediated Physcomitrella patens transformation

In the simplified transformation process for *Physcomitrella patens*, 5 mL of *Agrobacterium tumefaciens* strain EHA105 culture was directly added to each culture dish for 30 min treatment, the bacterial solution was removed using a sterile pipette and the culture plate was sealed with Parafilm. The gametophytes were then transferred to a 50-mL centrifuge tube. Subsequently, 20 mL of infection solution was added to the tube, after 30 min the mixture was returned to the culture plate. The culture plates were sealed and placed in the dark for 3 days. Following this, the cultures were transferred to standard *Physcomitrella patens* culture conditions for further growth. Validation was conducted one month after screening and culture to confirm the successful transformation of FBP in *Physcomitrella patens*.

### Plant material resources preparation

A total of 38 species were included in this study, categorized as 1 algae (*Chlamydomonas reinhardtii*), 2 bryophytes (*Physcomitrella patens, Marchantia polymorpha*), 1 lycophytes (*Diphasiastrum complanatum*), 1 ferns (*Ceratopteris richardii*), 3 gymnosperms (*Ginkgo biloba*, *Nageia nagi* and *Taxus chinensis*), 2 ANA basal angiosperms (*Amborella trichopoda, Nymphaea colorata*), 28 angiosperms (*Rhamnus hybrida*, *Rosa odorata*, *Solanum lycopersicum*, *Helianthus annuus*, *Rosa hybrida*, *Camellia japonica*, *Michelia figo*, *Freesia hybrida*, *Gossypium hirsutum*, *Gossypium barbadense*, *Prunus serrulate*, *Vigna radiata*, *Adenium obesum*, *Tulipa gesneriana*, *Begonia hiemalis*, *Phalaenopsis aphrodite*, *Dahlia pinnata*, *Syzygium grijsii*, *Dracaena sanderiana*, *Ranunculus asiaticus*, *Cyclamen persicum*, *Campanula medium*, *Senecio radicans*, *Bryophyllum pinnatum*, *Consolida ajacis*, *Nicotiana benthamiana*, *Catharanthus roseus*, *Ligustrum ovalifolium*). For the plants shown in Figure 3 specific germplasm resource are: *Adenium obesum* (Huanshanv), *Tulipa gesneriana* (Virgin), *Begonia hiemalis* (Huangse Ligehaitang), *Rosa hybrid* (Juicy Terrazza), *Phalaenopsis orchids* (Baitiane), *Helianthus annuus* (Vincent 2), *Gossypium barbadense* (Hai7124), *Vigna radiate* (Liaolv 27), *Dahlia pinnata* (Huanxiang). For the plants shown in the attached figure, they are all locally cultivated varieties from southern China.

### Database search

To identify candidate genes involved in the caffeic acid synthesis pathway, multiple database searches were conducted. Initially, the amino acid sequences of key genes (*4CL*, *C3’H*, *C3H*, *C4H*, *CSE*, *HCT*, *PAL*) were retrieved from the TAIR database (Barros *et al*., 2019). These sequences were then used as query sequences in BLASTP searches (E value < 1e^-2^) to identify additional candidate proteins in Table S1. The candidate genes were obtained from NCBI (https://www.ncbi.nlm.nih.gov/), Ginkgo DB (https://ginkgo.zju.edu.cn/) and Phytozome V13 (https://phytozome-next.jgi.doe.gov/) (Marchler-Bauer *et al*., 2015). Genes that did not show significant similarity to known genes involved in caffeic acid synthesis were removed from the candidate list based on the blast results and domain analysis from online database. This rigorous approach ensured that only relevant candidate genes were considered for further analysis in our study.

### Construction of the phylogenetic tree

The protein sequences of the seven genes involved in the caffeic acid synthesis pathway were aligned using MAFFT with default parameters (Katoh and Standley, 2013). Subsequently, phylogenetic trees for each gene were constructed separately using FastTree software, with 1000 resamples and the JTT model for maximum-likelihood analysis (Price, 2010). The accuracy of the phylogenetic trees was verified using the MEGA11 tools (Tamura *et al*., 2021). All amino acid sequences used for phylogenetic and alignment analysis in this resource study are provided in Table S1.

### *Agrobacterium*-mediated transient transformation in plants

Plasmids were transferred into *Agrobacterium tumefaciens* strain EHA105, which was cultured in LA Solid medium with 50 mg/L rifampicin and 50 mg/L kanamycin sulfate at 28°C for 48 h. The bacteria were then scraped and resuspended in a 500 mL washing solution containing 10 mM MgCl_2_ and 100 μM acetosyringone to reach an OD_600_ of 0.8-1.0. After centrifugation, the bacteria were resuspended in a liquid infiltration medium (1/4 MS salts, 1% sucrose, 100 μM acetosyringone, 0.005% Silwet L-77, pH 5.8) to the same OD_600_ range. The bacterial suspensions were incubated at room temperature without shaking for 2 h before use. Using a 1-mL plastic syringe, Agrobacterium solution was infiltrated into plant tissue after making a shallow puncture. The leaves were allowed to dry in the light for 1 h, followed by incubation in the dark at room temperature for 12 h. Transformed plants were then moved to the greenhouse and grown for an additional 2-3 days before sample observation. All cotton species were used for agroinfiltration at 20 days after seed germination, with the first true leaf. Mung bean plants, on the other hand, were injected on the first true leaves at 10 days after seed germination.

### Western blot analysis

Tobacco leaves transiently expressing fusion proteins were first homogenized in liquid nitrogen using a mortar and pestle. Each 0.1□g sample of homogenized leaves was then mixed with 200□μL of extraction buffer containing 50□mM Tris-HCl (pH 7.5), 150□mM NaCl, 0.5% TritonX-100, and Roche cocktail protease inhibitor. The mixture was centrifuged at 12000 g at 4°C for 10□min, after which the resulting supernatant was carefully transferred to a new tube. To this supernatant, the one-third volume of 4× Laemmli buffer was added, which consisted of 250□mM Tris-HCl (pH 6.8), 8% SDS, 40% glycerol, 4% β-mercaptoethanol, and 0.01% bromophenol blue. The mixture was then thoroughly mixed and denatured at 98°C for 10□min to obtain the denatured mixture containing the total proteins. Subsequently, the total proteins were separated by sodium dodecyl sulphate-polyacrylamide gel electrophoresis (SDS-PAGE) and transferred onto a polyvinylidene difluoride membrane (Millipore, USA). The membrane was then incubated with antibodies against YFP tags, while actin antibody was used as a loading control. The proteins were detected using ECL Western Blotting Detection Reagents (Sparkjade, China).

### LC-MS/MS analysis

The plant samples were harvested and immediately frozen in liquid nitrogen. They were then subjected to lyophilization in 50□mL Falcon tubes. Approximately 100□mg of the resulting lyophilized powder was measured and transferred to 5□mL extraction buffer, which contained 70% methanol. The extracts were then subjected to ultrasonication in a water bath for 30□minutes, followed by centrifugation at 13000□g for 15□min. The supernatants were filtered through a PVDF syringe filter with a pore size of 0.45□μm. The filtered extracts were transferred to glass vials for LC/MS analysis, as described in more detail previously (Zheng *et al*., 2023).

To conduct the SA analysis, leaves were harvested at the third day after agroinfiltration. SA was extracted using ethyl acetate spiked with labeled internal standards and analyzed using high-performance liquid chromatography combined with tandem mass spectrometry (HPLC/MS) system, following the previously described procedure (Lu *et al*., 2015). Each treatment was replicated three times.

### Quantitative PCR (qPCR)

To extract RNA, all leaves were initially flash frozen in liquid nitrogen and then homogenized using VeZol reagent (Vazyme Biotech, China). Subsequently, the 1□μg extracted RNA was utilized to synthesize first-stranded cDNA with the MonScriptTM RTIII Super Mix with dsDNase (Monad Biotech, China), following the manufacturer’s protocol. For qPCR analysis, gene transcript levels were determined using SYBR Premix and gene-specific primers on a LightCycler480 II Real-Time PCR machine (Roche, Switzerland). The PCR program included an initial step at 95°C for 1□min, followed by 40 cycles of 95°C for 10□s, 60°C for 20□s, and 72°C for 20□s. To ensure reliability, a minimum of three biological replicates were examined for each sample. The reference genes *NtEF1a, VrActin*, *GhUBQ7*, *HaEF1a*, *Ao5Srrna*, *TgUBQ10*, *PaUbi*, *BhACT2*, and *RhUbi* were employed for normalizing gene expression, and the primers utilized for amplification are detailed in Table S2.

### Subcellular localization

After the pCAMBIA1300-p35S-GhbHLH121-eGFP, pCAMBIA1300-p35S-NLS-mCherry and pCAMBIA1300-p35S-NahG-2A-P19 were transformed into the *Agrobacterium tumefaciens* strain EHA105, the resulting vectors were co-transfected into target tissuies. After culturing for 72□h at 28°C, fluorescence signals were observed and captured on a confocal microscope (Zeiss LSM880, Oberkochen, Germany).

### Luciferase complementation assay

*Agrobacterium tumefaciens* strain EHA105 containing the cLUC-GhbHLH121, GhPYE-nLUC, and NaP19 plasmids were co-injected into 20-day-old cotton leaves with the first true leaves. After two days, 0.15 mg/mL D-Luciferin solution was injected into the same site using a 1 mL plastic syringe. After a 20 min dark treatment, the leaves were imaged using a fully automated luminescence imaging system (Tanon5200, Shanghai, China).

### Plant imaging

The NIGHTSHADE LB985, a high-quality instrument from Germany, for imaging and calculating photon doses of plant bioluminescence signals. This instrument features a back-thinned ultra-sensitive CCD camera as its main component. During the experiments, the samples were placed inside a light-protected dark box at a height of approximately 40 cm from the top. Bioluminescent images were captured with a 60-second exposure time, and specific regions were selected for photon dose calculations. The data obtained was then exported for further analysis. After the bioluminescence measurements, ambient light images were taken using the same instrument settings. All other parameters were kept at their default values throughout the experiment.

### Statistical analyses

The quantitative data presented in this paper are expressed as mean□±□SD, obtained from a minimum of three biological replicates. GraphPad Prism 9 software was utilized for data visualization and plotting, while Excel was employed for statistical analysis. Statistical significance was determined using an unpaired two-tailed Student’s t-test for comparisons between two groups. The specific *P* values are indicated in the figures for reference.

## Acknowledgements

We thank Prof. Jian-Feng Li (MOE Key Laboratory of Gene Function and Regulation, School of Life Sciences, Sun Yat-sen University, Guangzhou, China) for sharing *Avrpto* and *AvrptoB* genes, Prof. Jun Zhao (Biotechnology Research Institute, National Key Facility for Crop Gene Resources and Genetic Improvement, Chinese Academy of Agricultural Sciences, Beijing, China) for providing the *ZmABP9* gene. We also thank Prof. Zhenghe Li (State Key Laboratory of Rice Biology, Institute of Biotechnology, Zhejiang University, Hangzhou, China) for providing the *P19* gene. This work was financially supported by the National Natural Science Foundation of China (32450506 and 32470288 to Hao Du), A.S. Mishin acknowledges financial support by Russian Science Foundation project number 24-74-10087.

## Author contributions

H.D. and T.W. initiated the project and designed the experiments, H.D., T.W., J.G., C.Q., H.C., X.G., Z.S., T.C., C.H., X.D. and L.L. performed experiments. T.W. and H.C. performed bioinformatics analyses. H.D. and T.W. wrote the manuscript. All authors discussed the results and commented on the manuscript.

