## Supplementary material for "Developing a versatile technology for agroinfiltration in multiple plants": supplemental data.docx

**Supplemental information**

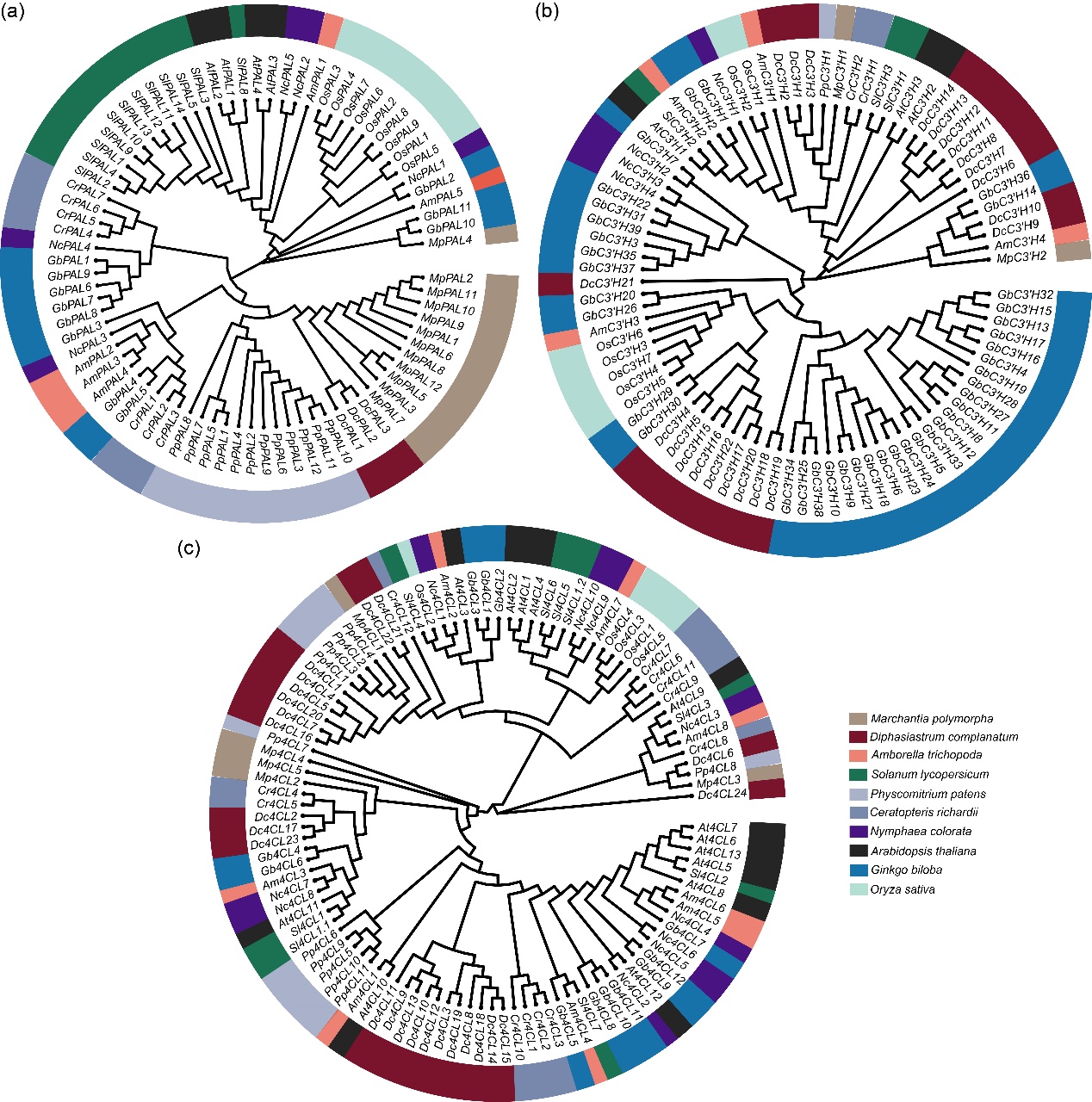

**Figure S1. Phylogenetic tree of PAL, C3’H and 4CL in plants orthologs.** The full-length amino acid sequence of PALs (a), C3’Hs (b) and 4CLs (c) from 10 plants species was retrieved with BLASTP (https://phytozome-next.jgi.doe.gov/, https://ginkgo.zju.edu.cn/), including *Marchantia polymorpha* (Map), *Physcomitrium patens* (Php), *Ginkgo biloba* (Gib), *Diphasiastrum complanatum* (Dic), *Ceratopteris richardii* (Cer), *Oryza sativa* (Ors), *Amborella trichopoda* (Amt), *Nymphaea colorata* (Nyc), *Solanum lycopersicum* (Sol) and *Arabidopsis thaliana* (Art). Sequence alignment was used to construct an NJ tree with MEGA 11 (Tamura *et al.*, 2021), using Poisson correction, pairwise deletion, and 1000 bootstrap replicates.

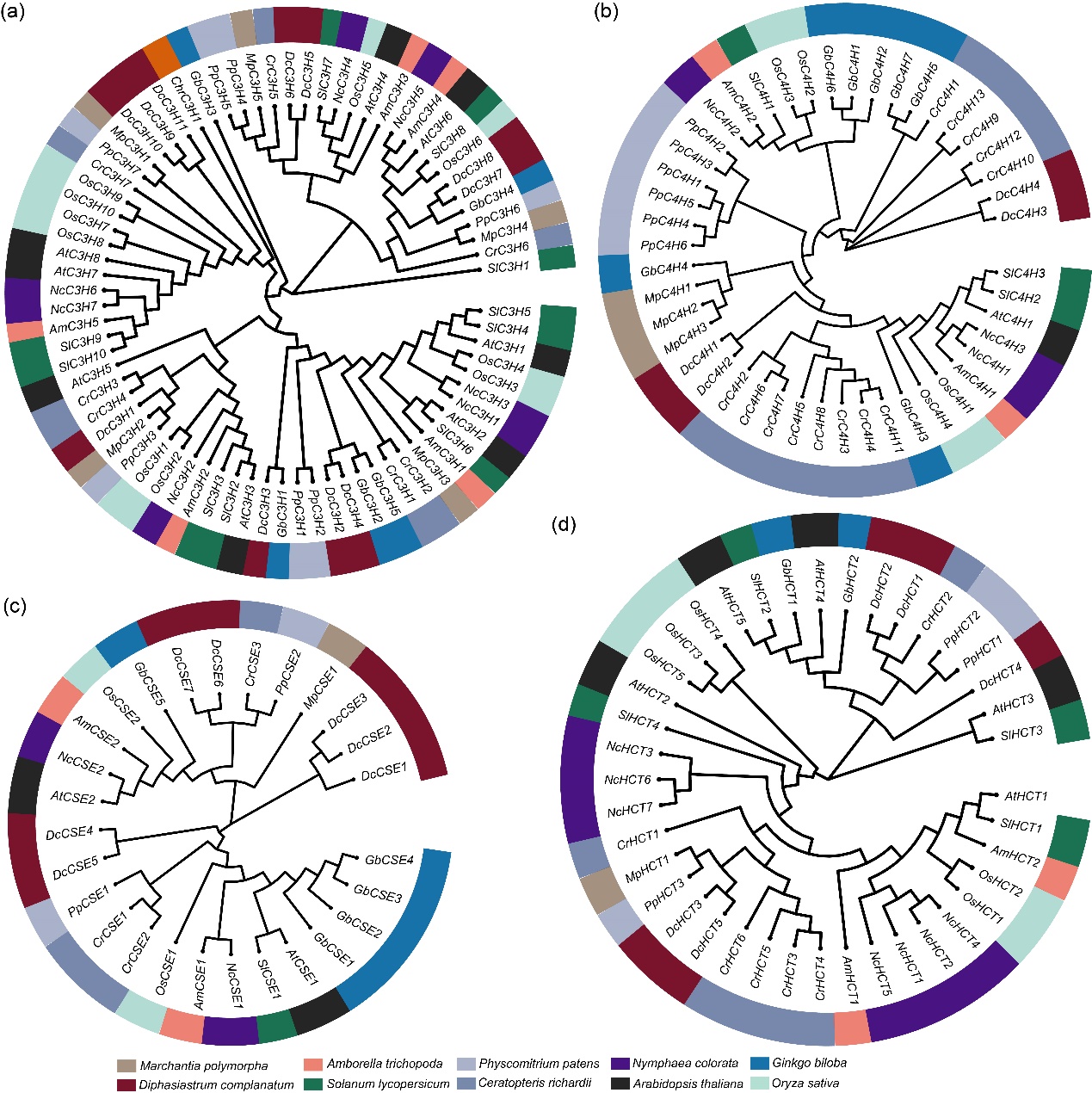

**Figure S2. Phylogenetic tree of C3H, C4H, CSE and HCT in plants orthologs.** The full-length amino acid sequence of C3Hs (a), C4Hs (b), CSEs (c) and HCTs (d) from 10 plants species was retrieved with BLASTP (https://phytozome-next.jgi.doe.gov/, https://ginkgo.zju.edu.cn/), including *Chlamydomonas reinhardtii* (Chr), *Marchantia polymorpha* (Map), *Physcomitrium patens* (Php), *Ginkgo biloba* (Gib), *Diphasiastrum complanatum* (Dic), *Ceratopteris richardii* (Cer), *Oryza sativa* (Ors), *Amborella trichopoda* (Amt), *Nymphaea colorata* (Nyc), *Solanum lycopersicum* (Sol) and *Arabidopsis thaliana* (Art). Sequence alignment was used to construct an NJ tree with MEGA 11(Tamura *et al.*, 2021), using Poisson correction, pairwise deletion, and 1000 bootstrap replicates.

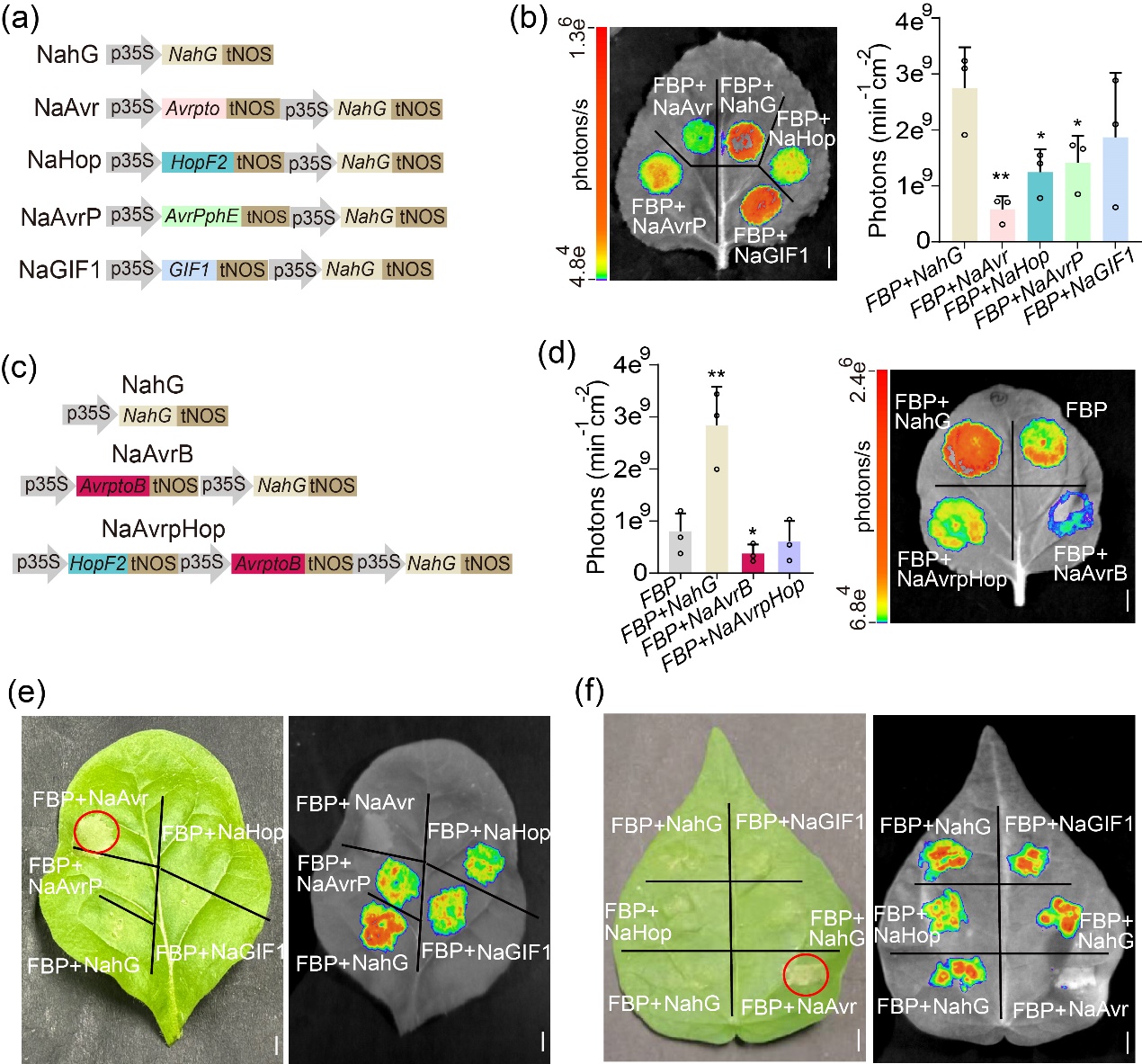

**Figure S3. The screen of enhanced effectors for agroinfiltration efficiency with *FBP* report system in multiple plants.** (a and b) Schematic illustrations (a) and agroinfiltration in *N. benthamiana* (b) with FBP report system. NahG (M60055), Avrpto (WP_011104823.1), HopF2 (AAK49537.1), AvrPphE (AAP23130.1), GIF1 (XP_037429183.1). Scale bars, 1 cm. Error bars indicate means ± SD (n = 3). Statistical significance was assessed using two-tailed Student’s *t*-tests (**P* ≤ 0.05, ***P* ≤ 0.01). (c and d) Schematic of agroinfiltration effectors (c) and agroinfiltration in *N. benthamiana* (d) with FBP-based reporter. Scale bars, 1 cm. Error bars indicate means ± SD (n = 3). AvrptoB (WP_011104378.1). Statistical significance was assessed using two-tailed Student’s *t*-tests (**P* ≤ 0.05, ***P* ≤ 0.01). (e and f) The screen of enhanced effector for agroinfiltration efficiency with FBP-based reporter in *N. tabacum* (e) and *Gossypium barbadense* (f). red circles indicate the necrotic spots. Scale bars, 1 cm.

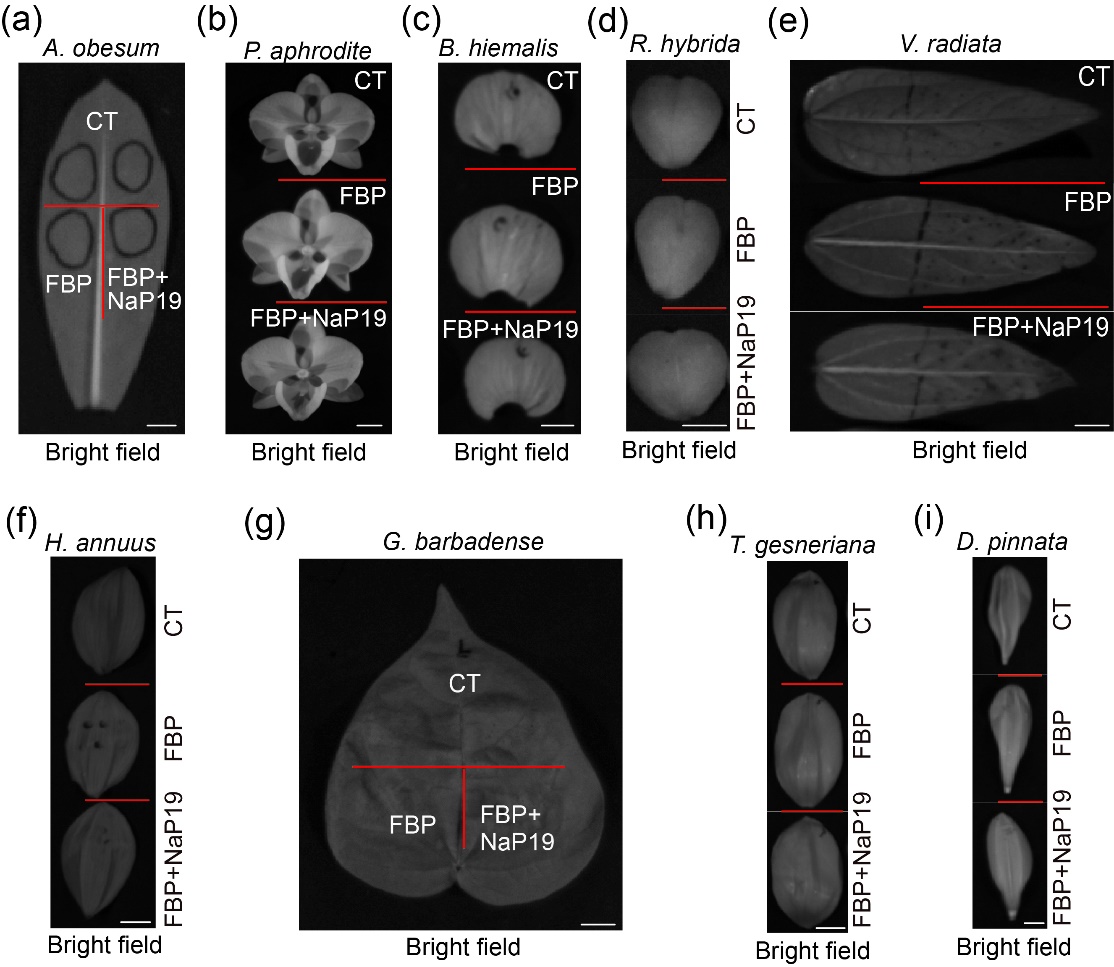

**Figure S4. The bright field of NaP19 enhances the agroinfiltration efficiency with the *FBP* report system in diverse plant species.** The bright field image of agroinfiltrated tissues in Figure 3a-i, *Adenium obesum* (a), *Phalaenopsis aphrodite* (b), *Begonia hiemalis* (c), *Rhamnus hybrida* (d), *Vigna radiate* (e), *Helianthus annuus* (f), *Gossypium barbadense* (g), *Tulipa gesneriana* (h), *Dahlia pinnata* (i), tissues agroinfiltrated with FBP and FBP+NaP19 modules respectively. Bioluminescent intensity analysis from agroinfiltrated leaves after 72 h. Scale bars, 1 cm.

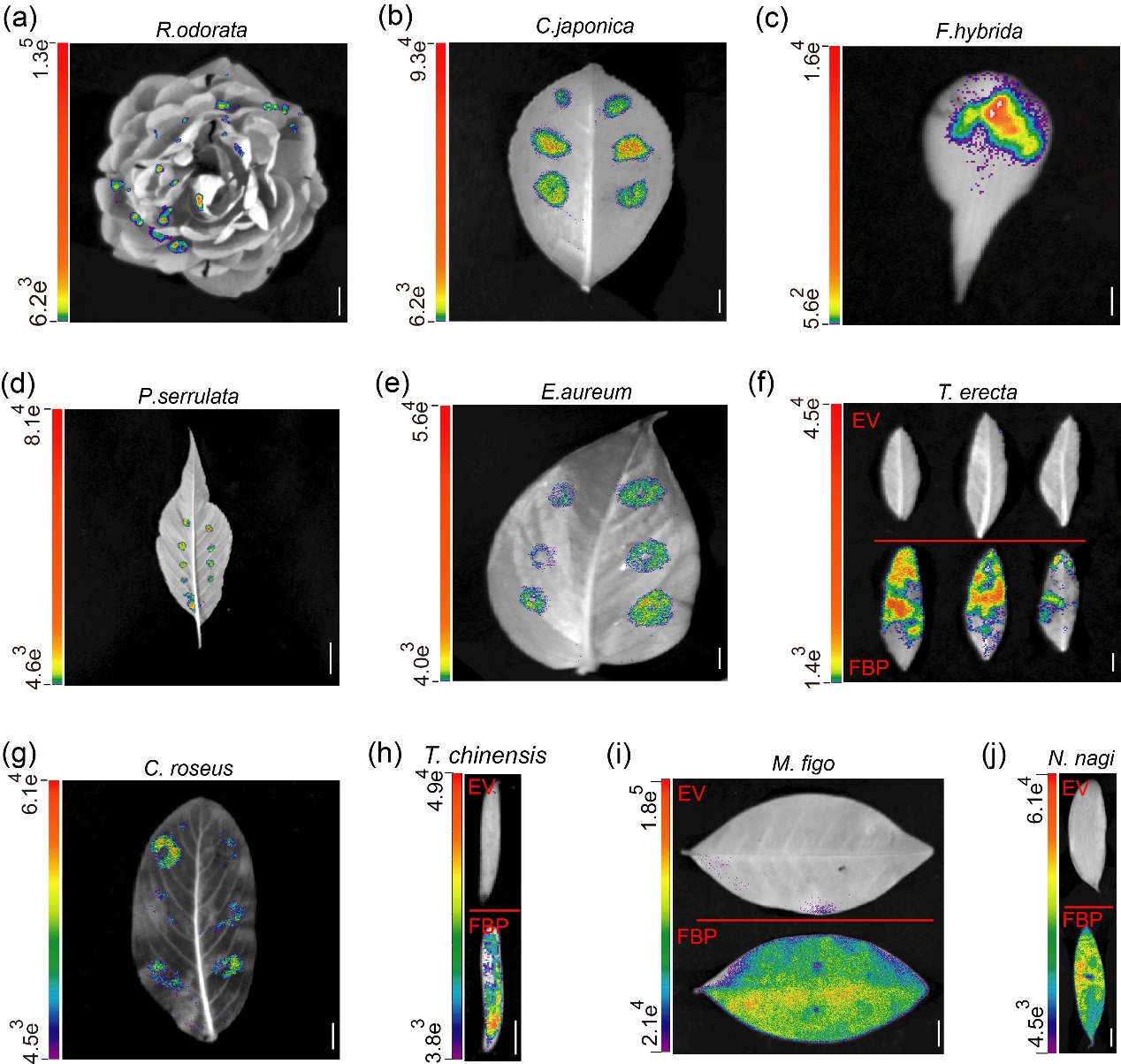

**Figure S5. The high-efficiency agroinfiltration of the tissues of multiple plant species.** (a-j) Optimized agroinfiltration of *Rosa odorata* (a), *Camellia japonica* (b), *Freesia hybrida* (c), *Prunus serrulata* (d), *Epipremnum aureum* (e), *Tagetes erecta* (f), *Catharanthus roseus* (g), *Taxus chinensis* (h), *Michelia figo* (i), *Nageia nagi* (j) tissues with FBP and EV (empty vector), respectively. Bioluminescent image captured by photographic instrument from agroinfiltrated tissues after 72 h. Scale bars, 1 cm.

**Table S1** Candidate protein sequence used in this study.

**Table S2** Primers used in this study.

**Table S3** Vectors information used in this study.
